# Deletion of *Dock7* Exons 3 and 4 Results in Reduced Trabecular Bone Microarchitecture

**DOI:** 10.64898/2025.12.31.696872

**Authors:** Talia Lizotte, Conner Lajoie, Sarah Porter, Sydney Lang, Felix A. Vesper, Daniel J. Brooks, Carlos Gartner, Matthew S. Alexander, Calvin Vary, Mary L. Bouxsein, Kathleen A. Becker

## Abstract

Dedicator of Cytokinesis 7 (DOCK7) has recently emerged as a regulator of skeletal homeostasis, but existing *Dock7* mutant models harbor only global mutations and are incompatible with tissue-specific deletion studies. We previously generated a *Dock7*-floxed allele in which exons 3–4 are flanked by LoxP sites. To validate the utility of this allele for future conditional strategies, we globally deleted exons 3–4 to generate *Dock7^em2/em2^* mice and characterized their skeletal phenotype. Homozygous *Dock7^em2/em2^* mice exhibited a diluted coat color and white belly spot, consistent with spontaneous *Dock7* mutations, whereas heterozygotes exhibited no spotting or coat changes. Bone microarchitecture was assessed in 21-week-old males and females. Global deletion of *Dock7* exons 3–4 resulted in a 30–37% reduction in trabecular bone volume in the distal femur and L5 vertebrae. Cortical bone thickness was unchanged, though sex-specific alterations in femoral area suggested altered appositional bone growth. Mass spectrometry (SWATH) analysis of *Dock7^em2/em2^* bone marrow stromal cells (BMSCs) identified DOCK7 protein levels similar to control mice, likely resulting from an alternative translational start site. Despite stable protein abundance prior to differentiation, BMSCs from *Dock7^em2/em2^* mice exhibited reduced mineralization and decreased *Bglap* expression, indicating attenuated osteoblast differentiation. These findings demonstrate that *Dock7* exons 3–4 are required for normal trabecular bone acquisition and osteoblast function. While the *Dock7-em2* allele likely encodes a truncated protein product, its recapitulation of the *Misty* skeletal phenotype confirms that exons 3–4 are essential for DOCK7 activity. These studies in the *Dock7^em2/em2^* mouse provide a foundation for future tissue-specific deletion studies utilizing the *Dock7* exon 3–4 deletion model to define the cellular roles of DOCK7 in regulating bone formation and trabecular architecture.

**Lay Summary:** Identifying genes that regulate bone mass is essential for understanding osteoporosis. Mutation of the *Dock7* gene in mice results in low bone mass, similar to human osteoporosis. Previous models could not isolate its role in specific tissues, limiting our ability to understand how DOCK7 controls bone mass. We developed a new mouse model where *Dock7* can be mutated in specific tissues. In this study, we bred mice to carry this mutation across all tissues, causing significant bone loss. This validates a flexible tool for future research to mutate DOCK7 in specific cell types and map its role in skeletal health.

## Introduction

Osteoporosis is a chronic skeletal disorder characterized by reduced bone mass and increased fracture risk, affecting approximately 54 million individuals in the United States [1]. The resulting fractures contribute substantially to morbidity and mortality, underscoring the need to better define the molecular pathways that regulate skeletal homeostasis. Bone integrity is maintained through a tightly regulated balance between osteoblast-mediated bone formation and osteoclast-driven bone resorption. Disruption of this equilibrium leads to skeletal fragility. In addition to local cellular mechanisms, bone remodeling is influenced by systemic factors including nervous system activity, adipose tissue function, and muscle mass. Increasing evidence suggests that Dedicator of Cytokinesis 7 (DOCK7) may represent a previously underappreciated regulator of these processes [2, 3].

Spontaneous mutation of DOCK7 in the *Misty* mice results in low bone mass, impaired trabecular microarchitecture, and reduced fat-free mass in both sexes [2, 3]. A reduction in cancellous bone is detectable as early as 8 weeks of age, suggesting an essential role for DOCK7 in bone acquisition [3]. Although early acceleration of bone loss was initially observed in female *Dock7* mutant mice between 8–16 weeks of age, subsequent analyses indicate that deficits are largely established by 16 weeks [2, 3]. Cortical expansion is impaired in female mutants, whereas long-term periosteal expansion in males has not been fully characterized [2]. Collectively, these findings support a primary role for DOCK7 in skeletal development and peak bone mass acquisition rather than progressive age-related bone loss.

At the cellular level, the loss of DOCK7 is associated with reduced osteoblast numbers and bone formation rates, concomitant with increased osteoclast numbers, suggesting that DOCK7 may influence both arms of bone remodeling [2, 3]. Additionally, systemic alterations, such as reduced brown adipose tissue thermogenesis and decreased lean mass, have been described, suggesting that DOCK7 regulates bone through both cell-autonomous and systemic mechanisms [3]. While these studies establish the *in vivo* importance of DOCK7, mechanistic investigations into cell-specific roles of DOCK7 have been limited by the available genetic models.

DOCK7 is a member of the Class C subgroup of DOCK180-related guanine nucleotide exchange factors (GEFs) [4]. It contains two conserved domains: Dock homology region 2 (DHR-2), which mediates GDP-to-GTP exchange on small GTPases such as Rac1 and Cdc42, and Dock homology region 1 (DHR-1), whose functions are less well characterized [5, 6]. Emerging evidence suggests that the DHR-1 domain participates in phosphoinositide signaling and may regulate AKT/mTORC pathways through TSC-dependent mechanisms [7–9]. Beyond its canonical GTPase-dependent functions, DOCK7 also modulates cellular processes, including migration, polarity, and differentiation, in diverse cell types through interactions with proteins such as TACC3 and ErbB4 [10–14]. Specifically, loss of DOCK7 reduces migration of bone marrow stromal cells, suggesting a direct role for DOCK7 in osteoblast migration [2]. Its role in neuronal polarity and Schwann cell function further highlights its broad biological significance [10, 15–18].

Reports of patients with compound heterozygous *DOCK7* mutations describe phenotypes including seizures, blindness, and craniofacial abnormalities with mutations within the N-terminus, TACC3 binding domain, and DHR2 domain, among other regions [19–21]. However, models for studying DOCK7 loss-of-function remain limited. RNAi-based approaches have been widely used for transient *DOCK7* ablation. However, these approaches are time-intensive and limited by efficiency, off-target effects, and lack of tissue specificity [10, 15]. Spontaneous *Dock7* mutant mouse models, such as *Misty* and *Moonlight*, have provided insights into DOCK7 function [2, 3, 12]. The *Misty* model, which has been the primary model for studying DOCK7 in bone up until this point, carries a premature stop codon in *Dock7* exon 18 and exhibits a diluted coat color and white belly spot [12]. DOCK7 was undetectable by western blotting in *Misty* mice, but in-depth analysis of DOCK7 protein products was not performed [3]. While these constitutive mutant models offer insights into the physiological role of DOCK7, they lack the flexibility for tissue-specific studies needed to understand DOCK7’s more nuanced role in disease and its potential therapeutic opportunities.

To overcome these limitations, a conditional *Dock7* floxed allele (*Dock7^fl/fl^*) was recently generated using CRISPR/Cas9 to flank exons 3 and 4 with LoxP sites [22]. Deletion of these exons is predicted to introduce a premature stop codon and recapitulate key features of spontaneous *Dock7* mutants. To determine whether this targeted deletion mimics the skeletal phenotype observed in other spontaneous *Dock7* mutants, the *Misty* mice, and therefore can be used as a reliable model to study DOCK7 in bone, we generated mice with homozygous global deletion of *Dock7* exons 3–4 (*Dock7^em2/em2^*). We assessed bone microarchitecture, body composition, and *in vitro* osteoblast differentiation in *Dock7^em2/em2^* mice and compared to *Dock7^+/+^* wild-type control mice. Our findings demonstrate that *Dock7^em2/em2^* mice exhibit reduced trabecular bone mass and impaired mineralization *in vitro*, similar to previously described DOCK7 loss-of-function models. Together, these findings show that the deletion of Dock7 exons 3–4 reproduces key skeletal features of established *Dock7* mutant models, supporting its utility for studying DOCK7-dependent regulation of bone and enabling future tissue-specific investigations.

## 2. Methods

### Mice and regulatory approval

The *Dock7-em1* allele (*Dock7* floxed allele, *Dock7-fl*), with exons 3 and 4 flanked by LoxP sites, was generated in the Mouse Genomic Modification Core at MaineHealth Institute for Research (MHIR), as previously described [22]. It was used to create the *Dock7-em2^Cjr^* (*Dock7-em2*) allele, with global deletion of *Dock7* exons 3–4. The *Dock7-em2* allele was used in these studies. Briefly, the *Dock7-em2* allele was generated by crossing the *C57BL/6J-Dock7^em1/em1^* mice with the *B6.Cg-Edil3^Tg(Sox2-cre)1Amc^/J* (*Sox2-cre*, The Jackson Laboratory, 8454) line in the laboratory of Dr. Clifford Rosen at MHIR. To eliminate the *Sox2-cre* allele, the *Dock7-em2* allele was bred away from the *Sox2-cre* by backcrossing to *CB57BL/6J* mice. Germline transmission of the *Dock7-em2* allele was confirmed in offspring. Mice were selected that harbored the *Dock7-em2* allele without the *Sox2-Cre* allele, and the *Dock7-em2* allele was bred to homozygosity (*Dock7^em2/em2^*) as confirmed by genotyping (S. Figure 1B). A schematic of the *Dock7^em2/em2^* generation is shown in Figure 1. Genotypes of the *Dock7 ^em2/em2^* mice were confirmed upon arrival at the University of New England. Experimental mice were generated by homozygous *Dock7^+/+^* or *Dock7^em2/em2^* breeding pairs within 2 generations of a heterozygous mating pairs. Transgenic mice were backcrossed to *C57BL/6J* mice after no more than 10 generations.

**Figure 1.**
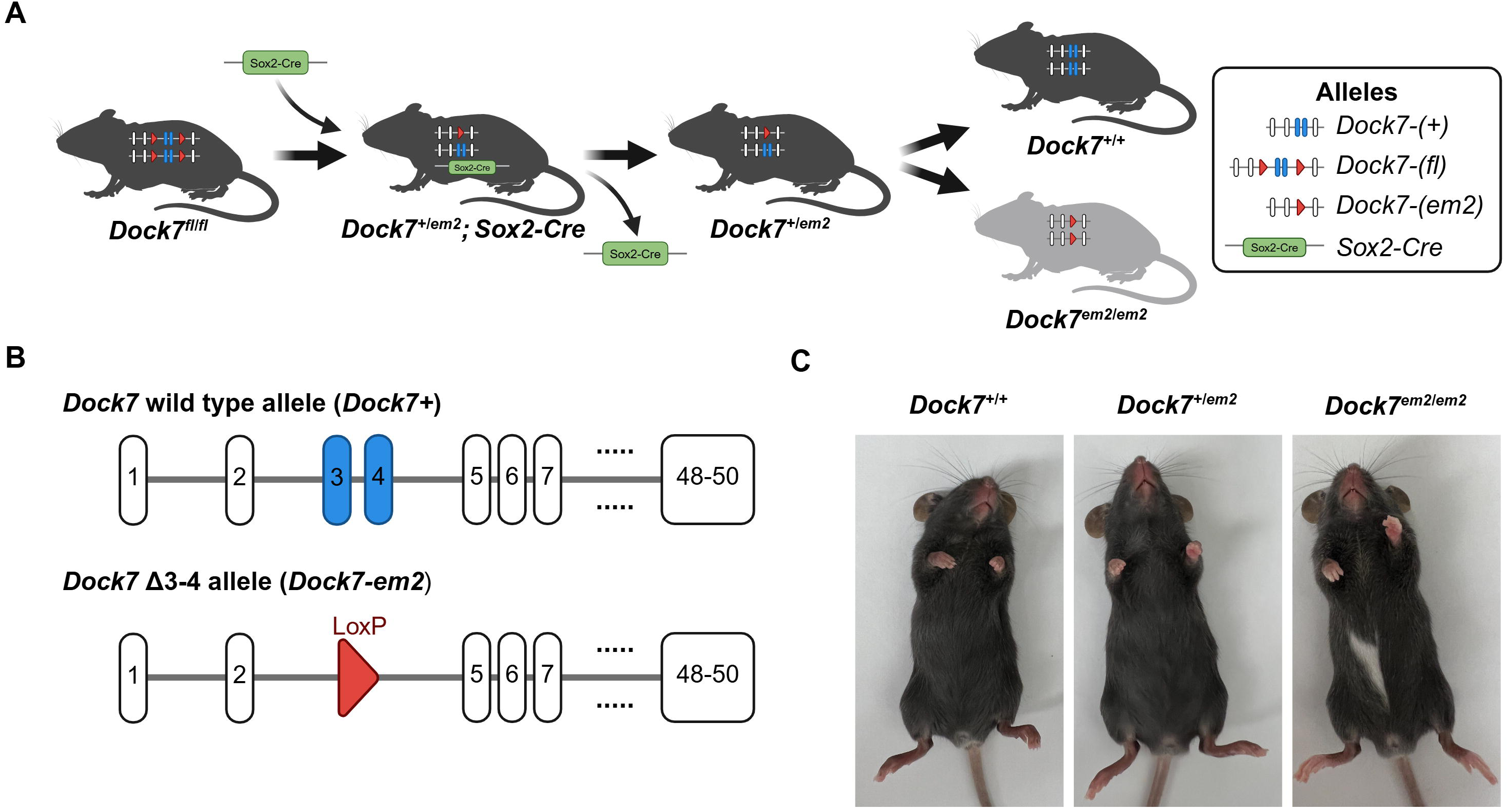
Deletion of *Dock7* exons 3–4 in the *Dock7^em2/em2^* mice results in a diluted coat color and white belly spot. (A) Targeted deletion of *Dock7* exons 3–4 was generated by crossing the *Dock7^em1/em1^* mice with *Sox2-cre* expressing mice. Mice harboring the *Dock7-em2* allele without the *Sox2-cre* transgene were selected, and the *Dock7-em2* allele was bred to homozygosity (*Dock7* ^em2/em2^). (B) Schematic of the *Dock7* wild type (+) and *Dock7* exons 3–4 deletion (*Dock7-em2)* transgenic allele. (C) Representative images of *Dock7^+/+^, Dock7^+/em2^, and Dock7^em2/em2^* mice. Images in (A) and (B) were created with BioRender.com. **Alt Text:** A three-part figure showing the genetic strategy and physical outcomes of *Dock7* gene deletion. Panel A is a workflow diagram showing the cross between *Dock7* floxed mice and *Sox2-Cre* mice to achieve a global deletion of *Dock7* exons 3 and 4. Panel B is a genomic map comparing the Dock7 wild-type allele to the mutant allele with deletion of *Dock7* exons 3 and 4. Panel C shows photographs of three mice: the wild-type and heterozygous mice display uniform dark coats, whereas the homozygous mutant mouse displays a distinctly lightened, diluted brown coat with a prominent white spot on its abdomen.

All animal studies followed National Institutes of Health (NIH) guidelines for the Care and Use of Laboratory Animals and were approved by the IACUC of the University of New England or MHIR. Bone phenotyping cohorts were housed and bred in the MHIR barrier vivarium, an AALAC-accredited facility. Mice were group housed on a 14:10-hour light/dark cycle and fed standard laboratory chow (Teklad irradiated global 18% protein diet, Envigo, 2918) with water ad libitum. Mice for *in vitro* differentiation of bone marrow stromal cells were group housed and bred at the University of New England, kept on a 12:12-hour light/dark cycle, and fed standard laboratory chow (Teklad global 18% protein diet, Envigo, 2018) with water ad libitum.

### Genotyping

Genotyping of the *Dock7-em2* and *Dock7+* alleles was performed on tail or toe clips incubated in 50 mM NaOH at 95°C for 1 hr and neutralized with 10% volume of 1M Tris, pH 8.0. DNA from tissue lysates was amplified using primers listed in S. Table 1, a Terra PCR Direct Polymerase Mix (Takara, 639270), and cycling conditions: 1. 98°C, 2 min; 2. 98°C, 10 sec; 3. 60°C, 15 sec; 4. 68°C, 1 min; 5. Steps 2-4, 26 cycles. The *Dock7+* (271 bp) and *Dock7-em2* (362 bp) PCR products were detected on a 3% agarose gel, using a 100 bp DNA ladder (New England Biolabs, N0467) as shown in S. Figure 1B.

**Table 1.** Deletion of *Dock7* exons 3–4 results in reduced femoral trabecular microarchitecture at 21 weeks of age.

|  | Female |  | Male |  | Two-way ANOVA *p*-values |  |  |
| --- | --- | --- | --- | --- | --- | --- | --- |
|  | *Dock7*<sup>+/+</sup><br>(n=9-10) | *Dock7*<sup>em2/em2</sup><br>(n=10) | *Dock7*<sup>+/+</sup><br>(n=9-10) | *Dock7*<sup>em2/em2</sup><br>(n=9-10) | Interaction | Sex | Genotype |
| **Femur length (mm)** | 15.2±0.1 | 14.9±0.1 | 15.2±0.1 | 15.0±0.1 | 0.9280 | 0.5805 | **0.0184** |
| **Tb.BV/TV (%)** | 5.12±0.26 | 3.24±0.25 | 19.40±1.42 | 12.19±1.63**** | **0.0197** | **<0.0001** | **0.0002** |
| **Tb.BMD (mgHA/cm<sup>3</sup>)** | 101.0±2.6 | 80.0±2.4 | 227.0±13.9 | 162.9±14.4*** | **0.0408** | **<0.0001** | **0.0002** |
| **Conn.D (1/mm<sup>3</sup>)** | 31.8±3.9 | 19.1±2.5 | 119.2±7.1 | 73.1±8.0**** | **0.0069** | **<0.0001** | **<0.0001** |
| **Tb.N (1/mm)** | 3.01±0.05 | 2.48±0.04*** | 4.47±0.13 | 3.75±0.10**** | 0.2776 | **<0.0001** | **<0.0001** |
| **Tb.Th (μm)** | 47.3±0.7 | 45.2±1.5 | 58.6±2.0 | 53.9±2.2 | 0.4604 | **<0.0001** | 0.0518 |
| **Tb.Sp (μm)** | 332±6 | 404±8**** | 210±8 | 258±7*** | 0.1149 | **<0.0001** | **<0.0001** |
| **SMI** | 3.35±0.08 | 3.39±0.08 | 1.62±0.13 | 2.30±0.17*** | **0.0110** | **<0.0001** | **0.0048** |

Data presented as mean±SEM.
Abbreviations: BMD, bone mineral density; BV, bone volume; Conn.D, connectivity density; N, number; pMOI, polar moments of inertia; Sp, separation; SMI, structural model index; Tb, trabecular; Th, thickness; TV, total volume.
Bold Values represent statistical significance with *p*<0.05. Statistical significance is indicated: \*, *p*<0.05; \*\*, *p*<0.01; \*\*\*, *p*<0.001; \*\*\*\*, *p*<0.0001.

### Bone and body composition phenotyping cohort analyses

#### General cohort information and tissue collection

All mice in the general phenotyping cohort were collected over a 4-month period. Male and female mice were grouped by genotyping and sample size was determine based on previous studies [2, 3]. *Dock7^+/+^* wild-type control mice (n=15, female; n=13 mice, male) and *Dock7^em2/em2^* experimental mice (n=15 mice/sex) were aged to 21 weeks and euthanized with carbon dioxide asphyxiation. No adverse events or unexpected health complications were observed during the study. Femora, tibiae, lumbar vertebrae, and adipose depots were fixed in 10% neutral buffered formalin (10% NBF) for 72 hrs and transferred to 70% EtOH for long-term storage.

#### Body composition and bone mineral density analysis by dual-energy x-ray absorptiometry (DXA)

DXA was performed on male and female 20-week-old *Dock7^+/+^* and *Dock7^em2/em2^* mice (n=14–15/sex/genotype) using the GE-Lunar PIXImus system. All mice underwent DXA analysis under isoflurane anesthesia to assess whole-body composition, excluding the head. Measurements included fat mass, total tissue mass, total areal BMD (aBMD), total areal BMC (aBMC), femoral areal BMD (fBMD) and femoral areal BMC (fBMC). Fat-free mass, which approximates lean mass, was calculated by subtracting fat mass from total mass. The central femoral diaphysis was further analyzed as a region of interest (ROI).

#### Micro-computed tomography (µCT) analysis

Bone microarchitecture analysis was performed according to established guidelines for µCT analysis of rodent bones [23]. Intact femora (n=10/group) and L5 vertebrae (n=10/group/sex) from *Dock7^+/+^* and *Dock7^em2/em2^* mice were randomly selected prior to analysis. Any bones found to have damage that impaired microarchitecture analysis were excluded. A benchtop µCT imaging system (µCT40, Scanco Medical AG, Brüttisellen, Switzerland) was used to scan bones at a 10µm^3^ isotropic voxel size, 70 kVP energy level, 114 µA intensity, and 200 ms integration time.

In femora, the trabecular bone microarchitecture was evaluated in a 1.5 mm (150 transverse slices) long region of the distal metaphysis beginning 200 µm superior to the peak of the growth plate and extending proximally. The trabecular region of the L5 vertebral body included the endocortical region beginning 100 µm inferior to the cranial endplate and extended to 100 µm superior to the caudal growth plate. Mineral density thresholds of 385 mgHA/cm^3^ and 370 mgHA/cm^3^ were used to segment bone from soft-tissue in the femur and L5 vertebrae, respectively. Trabecular microarchitecture was analyzed using the standard trabecular bone morphology script in the Scanco Evaluation Program. Cortical bone was assessed in 50 transverse µCT slices (500 μm long region) at the femoral mid-diaphysis, and the region included the entire outermost edge of the cortex. Cortical bone was segmented using a fixed threshold of 700 mgHA/cm^3^ and was analyzed using the Scanco Midshaft Evaluation script.

#### Adipose depot weights

Whole gonadal white adipose tissue (gWAT), inguinal white adipose tissue (iWAT), and interscapular brown adipose tissue (iBAT) depot weights from 21-week-old *Dock7^+/+^* and *Dock7^em2/em2^* mice (n=14–15/sex/genotype) were recorded following fixation in 10% NBF and then transferred to 70% EtOH. No sample weights were excluded from the analysis.

### Bone marrow stromal cell (BMSC) culture

#### BMSC isolation and general cell culture

Tibiae, femora, and iliac crests were isolated from age-matched male and female *Dock7^+/+^* and *Dock7^em2/em2^* mice at 6–9 weeks of age. Bone marrow was combined from 2 to 5 mice per genotype in each experiment. Total bone marrow was plated in MEMα (Thermo Fisher, 12571) supplemented with 10% fetal bovine serum (FBS, VWR, 97068-085, lot #164K18) and 1% penicillin/streptomycin (P/S) (Thermo Fisher, 15140122). After 48–72 hours, the non-adherent cell population was removed, and the adherent BMSCs were lifted using 0.25% trypsin EDTA (Sigma, T4049). BMSCs were plated at 1–2 million cells/well (1–2.1 × 10^5^ cells/cm^2^) in 6-well plates (Corning, 353046). All cultures were incubated at 37°C and 5% CO_2_ throughout the experiment.

#### BMSC osteogenic differentiation

Osteoblast differentiation was induced at 40–90% BMSC confluence. BMSCs were incubated osteoblast differentiation medium containing MEMα supplemented with 5% FBS, 1% P/S, 8.0 mM β-glycerol phosphate (Sigma, G9422), and 50 µg/mL ascorbic acid to induce osteogenic differentiation. Media was changed every 2–3 days.

#### Analysis of osteogenic differentiation by alkaline phosphatase and von Kossa staining

BMSCs from *Dock7^+/+^* and *Dock7^em2/em2^* mice were incubated for 7 days in osteogenic differentiation medium. On day 7 of differentiation, BMSCs were fixed in 4% paraformaldehyde (Electron Microscopy Sciences, 15710) and stained for alkaline phosphatase activity and mineralization. Alkaline phosphatase activity was detected using the Leukocyte Alkaline Phosphatase kit (Sigma, 86R-1KT). Mineral deposition was tested using von Kossa staining. Individual experiments included 3 wells/genotype and differentiation was repeated 4 times/sex with different animal cohorts. Wells exhibiting total detachment or visible peeling of the cell monolayer were prospectively excluded from the final analysis.

### Analysis of gene expression by real-time reverse transcription quantitative PCR (RT-qPCR)

#### RNA isolation and reverse transcription

Gene expression was analyzed in BMSCs isolated from *Dock7^+/+^* and *Dock7^em2/em2^* mice. Total RNA was isolated via standard extraction with Tri Reagent (Molecular Research Center, TR118) and resuspended in water. Samples were stored at -80°C for RNA analysis. cDNA was generated from 1 µg total RNA. RNA was treated with DNaseI and reverse transcribed using the High-Capacity cDNA Reverse Transcription kit (Thermo Fisher, 4368814) according to the manufacturer’s instructions. cDNA was diluted 1:5 prior to quantitative analysis.

#### Quantitative analysis of *Dock7* exons 3–4 deletion in BMSCs by RT-qPCR

Transcript expression of *Dock7* exons 3–4 was analyzed in BMSCs differentiated in osteogenic media for 7 days. cDNA was generated as described above. Experiments were repeated 3 times/sex with 3 wells/genotype. *Dock7* exons 3–4 levels were quantified using a TaqMan gene expression assay containing primers that span across *Dock7* exons 3–4 (Thermo Fisher, Mm01259842_g1, 4351372) and TaqMan Fast Advanced Master Mix (Thermo Fisher, 4444556). Samples were run on the Applied Biosystems QuantStudio3 System V.1.5.3 using cycle conditions: 1. 95°C, 20 sec; 2. 95°C, 1 sec; 3. 60°C, 20 sec; 4. Steps 2–3, 40 cycles. All target transcripts were normalized to the *Hprt* transcript reference assay (Thermo Fisher, Mm03024075_m1, 4331182). Replicates from all 3 experiments were combined for analysis, and no samples were excluded from analysis.

#### Quantitative analysis of osteoblast-related genes and additional *Dock7* exons by RT-qPCR

The mRNA expression of *Runx2*, *Bglap*, *Alpl*, *Dock7* exons 11–12, and *Dock7* exons 34–35 were analyzed in BMSCs differentiated in osteogenic media for 7 days. cDNA was generated as described above. Experiments were repeated 3 times/sex with 3 wells/genotype. Transcript expression was analyzed using AzuraQuant Green Fast qPCR HiRox Mix (Azura Genomics, AZ-2020) on the Applied Biosystems StepOnePlus System using cycle conditions: 1. 95°C, 3 min; 2. 95°C, 15 sec; 3. 60°C, 30 sec; 4. Steps 2–3, 40 cycles; 5. 55°C→95°C, 1°/sec; 6. 95°C, 15 sec. Sequences of target primers are provided in S. Table 1. All target transcripts were normalized to *Hprt* transcript levels. Replicates from all 3 experiments were combined for statistical analysis. Samples with indications of genomic DNA contamination or extreme deviations in *Hprt* expression were excluded from the analysis.

#### Mass spectrometry

BMSCs were isolated from individual male and female *Dock7^+/+^* and *Dock7^em2/em2^* mice (n=2 mice/group). BMSCs plated according to section 2.4 and harvested at 15–65% confluence. Briefly, cells were lifted with 0.25% trypsin EDTA, washed twice with PBS, snap frozen, and stored at -80°C prior to protein isolation. Samples from each mouse were separated into 1–3 replicates prior to proteomic analysis. As previously reported, tryptic peptides were separated by reverse phase liquid chromatography using a hand-fabricated C18 column [24]. Protein identification and quantification was accomplished using a SWATH mass spectrometry workflow on a SCIEX 5600 TripleTOF mass spectrometer [25]. Data were analyzed using the Peakview SWATH microapp and MarkerView software packages, as described previously [24]. Technical replicates (n=3) from all mice were combined prior to statistical analysis.

### Statistical analysis

Graphs are plotted as box-and-whisker plots, displaying median and sample range. Individual mice and cell culture replicates are displayed as units of analysis. All statistical analyses were performed using GraphPad Prism version 11.0.3 (GraphPad Software, Boston, MA, USA). Statistical significance was determined using two-way ANOVA with Šidák’s multiple comparisons post-hoc test or Student’s T-test where appropriate. Statistical significance is indicated: *, *p*<0.05; **,*p*<0.01; ***, *p*<0.001; ****, *p*<0.0001.

### Graphic representation

Graphic figures were generated using GraphPad Prism 11.0.3 (GraphPad Software, Boston, MA, USA) and modified in Adobe Illustrator 2025. Diagrams were created in BioRender.

## Results

### Generation of the *Dock7-em2* allele

A *Dock7* allele with a deletion of exons 3–4 (*Dock7-em2*) was generated from the *Dock7^em1/em1^* mouse (*Dock7^fl/fl^*, Figure 1AB). This deletion is predicted to create a premature stop codon in the *Dock7* mRNA transcript (S. Figure 1A). Homozygous deletion of *Dock7* exons 3–4 in mice (*Dock7^em2/em2^*) was confirmed through genotyping (S. Figure 1B). *Dock7^em2/em2^* mice exhibited a diluted coat color and white belly spot, consistent with previously reported phenotypes in *Misty*, Moonlight, and CRISPR-mediated *Dock7* exon 3–4 deletion models [12, 22, 26]. Heterozygous *Dock7^+/em2^* mice did not display coat color changes (Figure 1C).

### Molecular analysis of the *Dock7-em2* allele in bone marrow stromal cells (BMSCs)

To demonstrate the impact of deletion of *Dock7* exons 3–4 at the *Dock7* protein level, protein expression for *Dock7* was quantified using tandem mass spectrometry in BMSCs isolated from *Dock7^+/+^* and *Dock7^em2/em2^* mice prior to osteogenic differentiation. High-confidence peptides associated with the DOCK7 protein were identified by sequential window acquisition of all theoretical mass spectra (SWATH) workflows in protein samples from both genotypes and used to quantify DOCK7 protein levels (S. Figure 2). These data indicate the presence of DOCK7 protein in BMSCs isolated from both *Dock7^+/+^* and *Dock7^em2/em2^* mice. DOCK7 protein abundance was not significantly different between genotypes (*p =* 0.76) as determined by two-way ANOVA analysis. However, DOCK7 protein expression was reduced in BMSCs from male mice when compared to BMSCs isolated from female mice (*p =* 0.0007) (Figure 2). High-confidence peptides were detected in the DHR2 domain in lysates from BMSCs of both genotypes (S. Figure 2). Since high-confidence DOCK7 peptides were not detected in *Dock7* exons 1–4, the loss of the N-terminal peptides prior to the exon 3–4 deletion or peptides within exons 3–4 could not be evaluated. A low-to-medium-confidence peptide was detected in *Dock7* exon 4 in both genotypes (S. Figure 2). Because both genotyping (S. Figure 1) and transcript analysis confirmed that this region was not detected (Figure 4) and that the peptide exhibits sequence homology with other mouse proteins, the evidence does not support an association of this peptide with DOCK7. Spectra and sequencing data are shown for representative high-confidence DOCK7 peptides (S. Figures 3–4).

**Figure 2.**
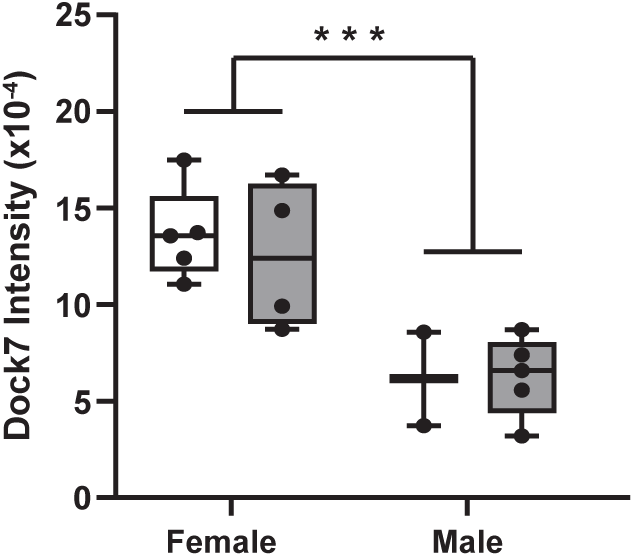
BMSCs from *Dock7^em2/em2^*mice had similar levels of DOCK7 protein compared *Dock7^+/+^* mice. Quantitative protein analysis of DOCK7 was performed by mass spectrometry in BMSCs isolated from *Dock7^+/+^* and *Dock7^em2/em2^* mice. Points represent individual replicates from all experiments. Data was analyzed by two-way ANOVA with Šídák’s test for post hoc analysis. Statistical significance is indicated: *, *p*<0.05; **,*p*<0.01; ***, *p*<0.001; ****, *p*<0.0001. **Alt text**: A one-panel molecular validation dataset of the *Dock7* exons 3–4 deletion in primary mouse bone marrow stromal cells (BMSCs). The panel contains a bar graph of DOCK7 protein abundance quantified by SWATH mass spectrometry. The graph shows similar DOCK7 protein levels between genotypes, although male BMSCs express significantly less total DOCK7 protein than female BMSCs, regardless of genotype.

### Bone microarchitecture analysis

To evaluate the effect of the *Dock7* exons 3–4 deletion on bone microarchitecture, femora and L5 vertebrae from male and female *Dock7^em2/em2^* mice at 21 weeks of age were analyzed and compared to age- and sex-matched *Dock7^+/+^* control mice. Two-way ANOVA demonstrated a robust reduction in Tb.BV/TV in *Dock7^em2/em2^* mice (*p =* 0.0002) (Table 1). Although post-hoc pairwise comparisons revealed a more pronounced, statistical deficit in male mice (*p* < 0.0001), the mean percentage reduction was identical (37%) between both sexes, suggesting a consistent biological impact of the mutation regardless of baseline sex differences in bone volume. Changes in femoral cancellous bone parameters, such as Conn. D., Tb.N, Tb.Th, and Tb.Sp were consistent with reduced trabecular bone in the distal femora with deletion of *Dock7* exons 3–4. Similarly, Tb.BV/TV was also significantly reduced in the L5 vertebrae in Dock7^em2/em2^ mice (*p* < 0.0001), with post-hoc pairwise comparisons revealing a similar deficit in both female (*p* <0.0001, 37%) and male (*p* < 0.0001, 30%) mice (Table 3). Changes in other cancellous bone parameters were consistent with reduced trabecular bone in the L5 vertebrae with deletion of *Dock7* exons 3–4.

No significant changes in cortical thickness (Ct.Th) were observed in femora by genotype (*p* = 0.56) as detected by two-way ANOVA. Other cortical parameters such as total area (Tt.Ar, *p* = 0.17), cortical area (Ct.Ar, *p* = 0.08), and medullary area (Ma.Ar, p = 0.42) did not meet the threshold for statistical significance due to sex-specific differences in cortical phenotype (Table 2). In femora from female *Dock7^em2/em2^* mice, increases were present for Tt.Ar (*p* = 0.007, 8.3%), Ct.Ar (*p* = 0.38, 2.5%), and Ma.Ar (p=0.0004, 14%) as detected by Student’s T-test. In femora from male *Dock7^em2/em2^* mice, decreases were present for Tt.Ar (*p* = 0.004, 12%), Ct.Ar (*p* = 0.02, 11%), and Ma.Ar (p=0.006, 13%) as detected by Student’s T-test. These findings indicate that deletion of *Dock7* exons 3–4 may result in altered appositional bone growth, with distinct effects between sexes.

**Table 2.**
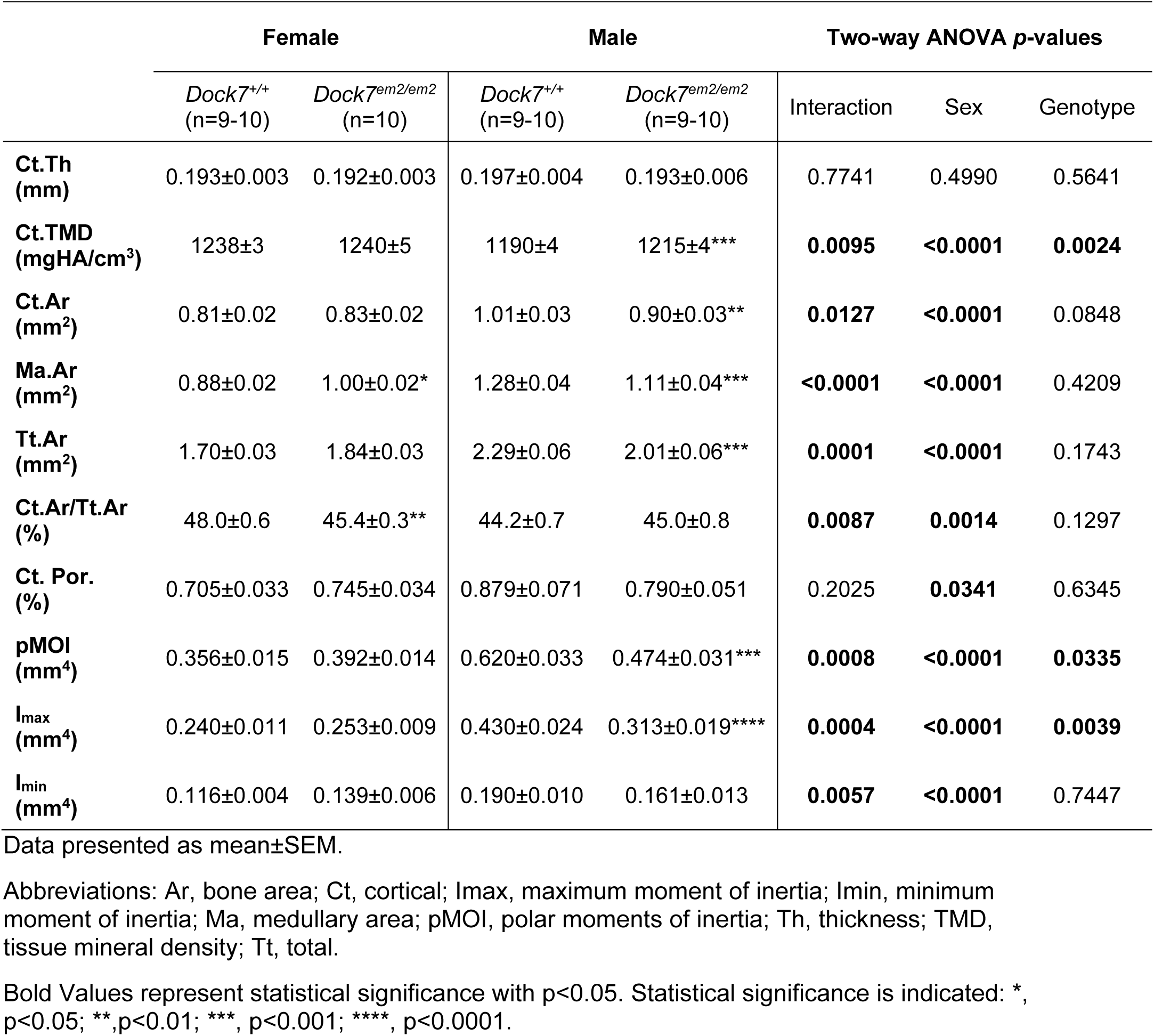
Deletion of *Dock7* exons 3–4 did not result in a detectable shift in cortical thickness at 21 weeks of age.

**Table 3.** Deletion of *Dock7* exons 3–4 results in reduced trabecular microarchitecture in the L5 vertebrae at 21 weeks of age.

|  | Female |  | Male |  | Two-way ANOVA *p*-values |  |  |
| --- | --- | --- | --- | --- | --- | --- | --- |
|  | *Dock7*<sup>+/+</sup><br>(n=9-10) | *Dock7*<sup>em2/em2</sup><br>(n=10) | *Dock7*<sup>+/+</sup><br>(n=9-10) | *Dock7*<sup>em2/em2</sup><br>(n=9-10) | Interaction | Sex | Genotype |
| **BV/TV (%)** | 21.40±0.50 | 13.44±0.85**** | 25.64±0.88 | 17.83±0.74**** | 0.9285 | **<0.0001** | **<0.0001** |
| **Conn.D (1/mm<sup>3</sup>)** | 92.1±5.1 | 69.2±6.6* | 156.6±6.4 | 96.6±7.0**** | **0.0062** | **<0.0001** | **<0.0001** |
| **Tb.N (1/mm)** | 3.60±0.10 | 2.95±0.12*** | 4.77±0.10 | 3.72±0.14**** | 0.1021 | **<0.0001** | **<0.0001** |
| **Tb.Th (μm)** | 58.5±0.7 | 53.8±0.8** | 55.4±1.8 | 52.5±0.8 | 0.4050 | 0.0549 | **0.0013** |
| **Tb.Sp (μm)** | 277±9 | 339±13** | 201±5 | 267±16*** | 0.8637 | **<0.0001** | **<0.0001** |
| **SMI** | 0.71±0.04 | 1.47±0.08**** | 0.61±0.08 | 1.15±0.05**** | 0.0917 | **0.0025** | **<0.0001** |

Data presented as mean±SEM.
Abbreviations: BV, bone volume; Conn.D, connectivity density; N, number; Sp, separation; SMI, structural model index; Tb, trabecular; Th, thickness; TV, total volume.
Bold Values represent statistical significance with *p*<0.05. Statistical significance is indicated: \*, *p*<0.05; \*\*, *p*<0.01; \*\*\*, *p*<0.001; \*\*\*\*, *p*<0.0001.

### Body composition and bone mineral density

Body composition was analyzed by DXA analysis at 20 weeks of age in male and female *Dock7^em2/em2^* mice and compared to age- and sex-matched *Dock7^+/+^* control mice. Two-way ANOVA demonstrated a reduction in total body weight in *Dock7^em2/em2^* mice (*p =* 0.0004) (Figure 3). The post-hoc pairwise comparisons revealed a statistical deficit in male mice (*p =* 0.0001, 13%) with a lower numerical decrease in female mice (*p =* 0.58, 3.8%). A decrease in fat-free mass was present in *Dock7^em2/em2^* mice (*p*<0.0001) by two-way ANOVA (Figure 3). The post-hoc pairwise comparisons revealed a statistically significant reduction in male mice (p<0.0001, 14%) and a numerical decrease in female mice (*p =* 0.18, 4.6%), similar to the decreases found in total body mass. Variations within fat-free mass, as calculated as a percentage of body weight, were not identified (*p =* 0.51). Changes in fat mass between genotypes were not detected by two-way ANOVA, whether calculated from direct measurements (*p =* 0.73) or as a percentage of total body mass (*p =* 0.51) (Figure 3). Ex vivo analysis revealed a consistent numerical reduction in two-way ANOVA across all adipose depots analyzed, including inguinal (*p =* 0.11), gonadal (*p =* 0.11), and brown (*p =* 0.083) fat pads. However, the dataset exhibits too much variance to establish a definitive statistical effect (S. Figure 5). A reduction in total (*p*<0.0001) and femoral (*p =* 0.0026) aBMD and total (*p*<0.0001) and femoral (*p*<0.0006) aBMC were detected by two-way ANOVA in *Dock7^em2/em2^* mice (S. Figure 6). As reductions in total body weight and fat-free mass are similar between sexes, these results suggest that the lower body weight observed in the *Dock7^em2/em2^* mice is likely a direct result of lower fat-free mass, with changes in fat mass being a minor contributor.

**Figure 3.**
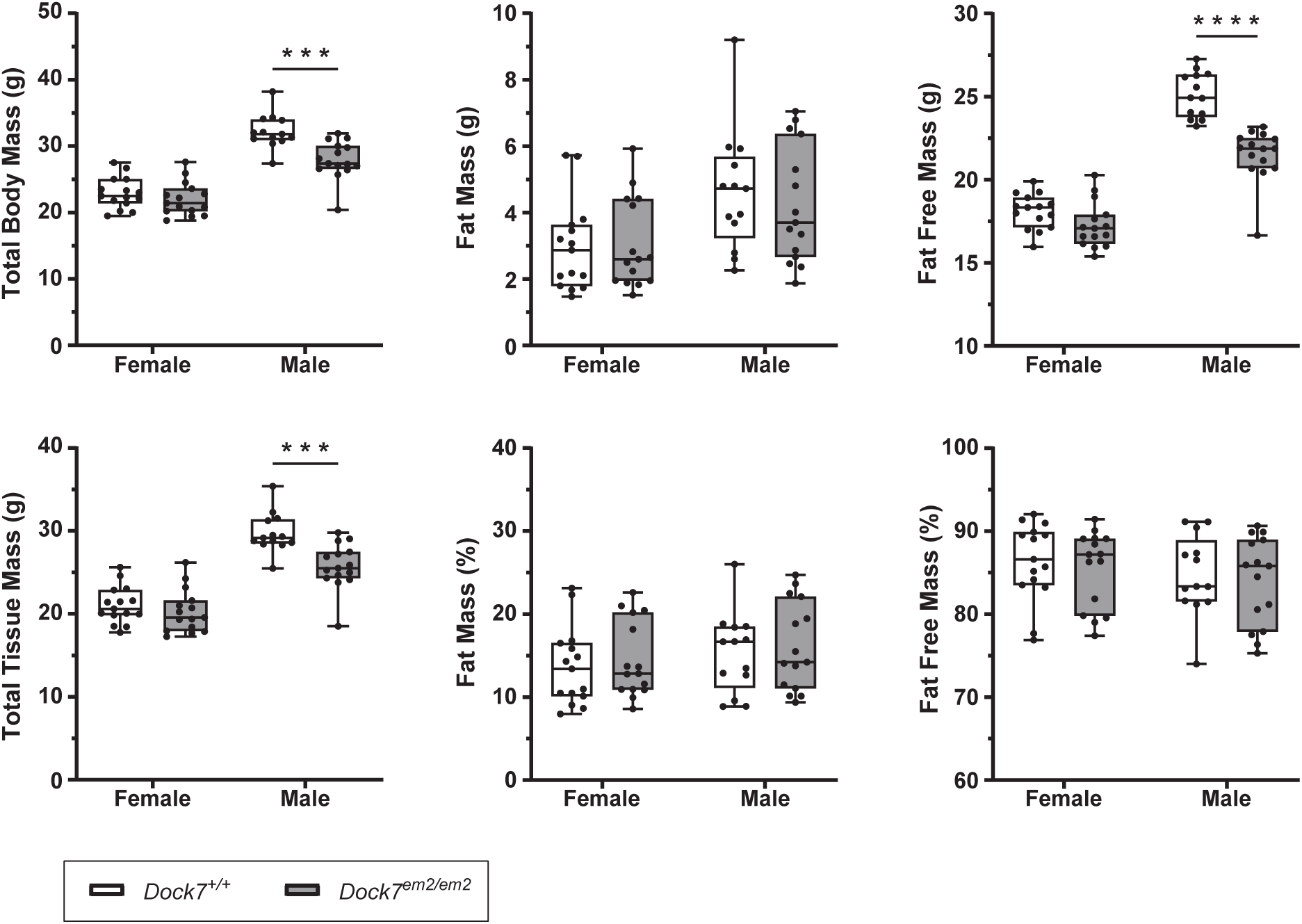
Body composition in *Dock7^em2/em2^* mice at 20 weeks of age. Body composition was measured by DXA analysis in male and female *Dock7*^em2/em2^ mice and compared to *Dock7^+/+^* control mice (N=14–15 mice/group). Points represent each mouse. Data was analyzed by two-way ANOVA and Šídák’s test for post-hoc analysis. Statistical significance is indicated: *, *p*<0.05; **,*p*<0.01; ***, *p*<0.001; ****, *p*<0.0001. **Alt text:** Six bar graphs illustrating DXA body composition in male and female *Dock7* wild-type vs. *Dock7* mutant mice at 20 weeks of age. Graphs show total body mass, fat mass, fat-free mass, and total tissue mass in grams. Fat mass and fat-free mass are also calculated as a percentage of total tissue mass as well. Decreases in total body mass, total tissue mass, and fat-free mass was present in both sexes. Equivalent levels of fat mass in grams as well of fat mass and fat-free mass as expressed as a percentage of total tissue mass were presented.

### *In vitro* osteogenic differentiation

The role of DOCK7 in osteogenic differentiation was explored using bone marrow stromal cells (BMSCs) isolated from *Dock7^em2/em2^* and *Dock^7+/+^* mice. Osteogenic differentiation was stimulated *in vitro* and analyzed at Day 7 for mineral deposition. BMSCs isolated from *Dock7^em2/em2^* mice showed a decrease in mineralization by von Kossa staining (Figure 4A). *Runx2* gene expression remained statistically equivalent following mutation of *Dock7* (female, *p =* 0.75; male, *p =* 0.12). In contrast, *Alpl* expression displayed a sex-specific pattern, characterized by a clear downregulation in male-derived BMSCs (*p =* 0.004) that was absent in female-derived cells (*p =* 0.77) as detected by Student’s t-test. Expression of the late-osteoblast gene, *Bglap,* was decreased in BMSCs from female (*p =* 0.03) and male (*p =* 0.0001) *Dock7^em2/em2^* mice (Figure 4BC). Lastly, Dock7 mRNA levels were analyzed at the exon 3–4, 11–12 and 34–35 boundaries to determine the impact of the *Dock7* exons 3–4 deletion on overall transcript levels. Dock7 exons 3–4 were undetectable in both sexes, confirming deletion of the exons. BMSCs from female *Dock7^em2/em2^* mice had a 58% (*p =* 0.0003) and 56% (*p*<0.0001) reduction at both loci, respectively. BMSCs from male *Dock7^em2/em2^* mice had a 71% (*p =* 0.0001) and 73% (*p*<0.0002) reduction at both loci, respectively (Figure 4BC). These results demonstrate that deletion of *Dock7* exons 3–4 reduced DOCK7 transcript levels during osteogenic differentiation and impaired *in vitro* osteoblast differentiation.

**Figure 4.**
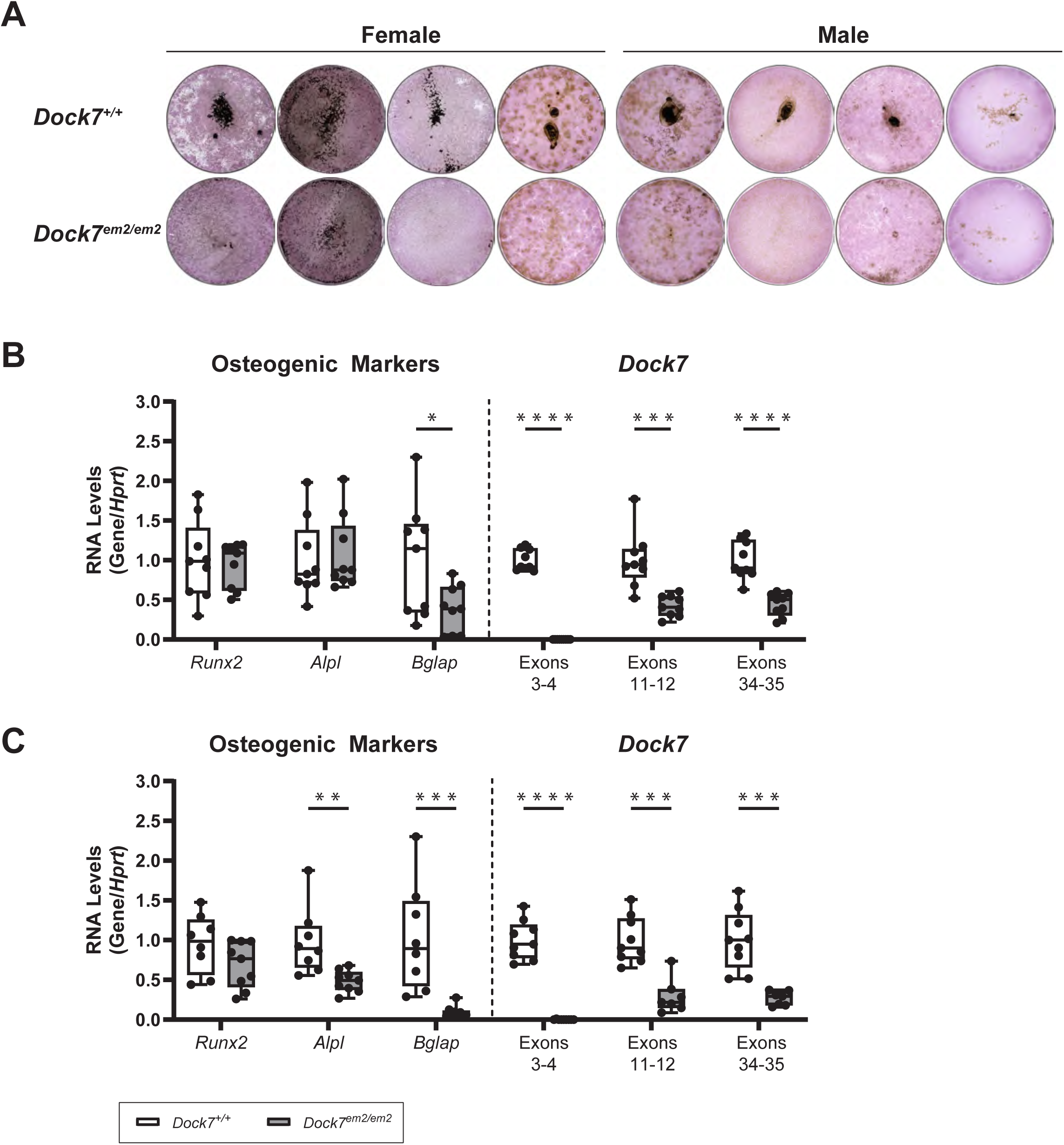
Deletion of *Dock7* exons 3–4 impaired *in vitro* mineralization and reduced *Bglap* gene expression in BMSCs. Osteogenic differentiation was assessed in BMSCs that were isolated from *Dock7^+/+^* and *Dock7^em2/em2^* mice. (A) Representative image of von Kossa and alkaline phosphatase staining after 7 days in osteogenic media. Representative wells are shown from 4 experiments. Expression of osteoblast-related genes (*Runx2*, *Alpl*, and *Bglap) and Dock7* (exons 3–4, 11–12, and 34–35) was analyzed in BMSCs from female (B) and male (C) mice at day 7 of osteogenic differentiation. Points represent individual replicates for all experiments. Data was analyzed by Student’s t-test within each gene. Statistical significance is indicated: *, *p*<0.05; **,*p*<0.01; ***, *p*<0.001; ****, *p*<0.0001. **Alt text:** Three-panel figure of osteoblast differentiation in primary BMSCs. Panel A shows culture wells at day 7 of differentiation; *Dock7* mutant wells demonstrate a reduction in dark-stained mineralized matrix via von Kossa staining compared to wild-type controls. Panels B (female) and C (male) display qPCR gene expression at day 7 of osteogenic differentiation. Results show significant downregulation of *Bglap* mRNA in both sexes. There is a male-specific downregulation of *Alpl* mRNA in *Dock7* mutant cells. *Runx2* mRNA expression remains statistically unchanged in female and male mice. In both sexes, *Dock7* mRNA shows an absence of signal within exons 3–4 and a significant reduction in the exons 11–12 and 34–35 in mutant cells.

## Discussion

These studies characterize a bone phenotype in a new model of *Dock7* mutation. Genomic deletion of exons 3–4 resulted in similar levels of DOCK7 protein in the *Dock7^em2/em2^* and *Dock7^+/+^* mice as assessed by mass spectrometry. The deletion of *Dock7* exons 3–4 resulted in a reduction in trabecular bone volume fraction (Tb.BV/TV) in the femur and L5 vertebrae across both sexes. The low trabecular bone volume observed in *Dock7^em2/em2^* mice was accompanied by a reduced expression of late osteoblast gene markers in BMSCs differentiated from *Dock7^em2/em2^* mice, highlighting the functional consequences of deletion of *Dock7* exons 3–4 in the *Dock7-em2* allele. With reduced levels of *Dock7* transcript in BMSCs following osteogenic differentiation, our data suggest that *Dock7* exons 3–4 may be necessary for *Dock7* transcript stability and protein function.

The observed trabecular bone phenotype in *Dock7^em2/em2^* mice aligns with findings from previous studies using *Misty* mice, a model with no detectable DOCK7 protein [3]. In *Misty* mice, reductions in Tb.BV/TV ranged from 22–60% at 16 weeks of age [2, 3]. Consistent with this, *Dock7^em2/em2^* mice exhibited trabecular bone loss within the narrower range of 30–37%. The similar degree of cancellous bone loss between these models supports the conclusion that DOCK7 is critical for maintaining approximately one-third of the trabecular bone mass in the femur and vertebrae. Together, these results validate *Dock7^em2/em2^* mice as a robust and reproducible model for studying the role of DOCK7 in cancellous bone.

Cortical bone phenotypes in *Dock7^em2/em2^* mice revealed sex-specific differences, consistent with prior studies of *Misty* mice [2, 3]. At 21 weeks, *Dock7^em2/em2^* mice had statistically equivocal Ct.Th compared to control mice. Distinct changes in Tt.Ar, Ct.Ar, and Ma.Ar. demonstrated an increase in appositional bone growth in female *Dock7^em2/em2^* mice while male mice showed a reduction in apposition bone growth. This sex-dimorphic pattern aligns with findings from *Misty* mice, where female mice displayed early increases in periosteal and endosteal circumference followed by slow appositional bone growth. Over time, female *Misty* mice also developed reductions in Ct.Th, a phenotype not observed in males [2, 3]. However, because Ct.Th appeared similar across genotypes in the *Dock7^em2/em2^* cohort, the deletion of exons 3–4 may present a more subtle cortical phenotype than that observed in the *Misty* mice, highlighting a potential difference in model-specific presentation. While a direct comparison is complicated by differences in age between this study and prior reports, these data suggest that DOCK7 may play a sexually dimorphic role in regulating cortical bone growth and maintenance. Further studies utilizing a longitudinal analysis of cortical bone, rather than the snapshot provided in this analysis, are needed to fully evaluate changes in apposition bone growth with deletion of *Dock7* exons 3–4, and compare to previous studies.

Deletion of *Dock7* exons 3–4 also impaired *in vitro* osteoblast differentiation with a reduction in *in vitro* mineralization and diminished expression of late osteoblast-gene markers in *Dock7^em2/em2^* mice, particularly in BMSCs from female mice. These results parallel observations in female *Misty* mice, where *Dock7* mutations reduced *Alpl* and *Bglap* expression during calvarial osteoblast differentiation [3]. However, this study expands upon prior work by demonstrating that *in vitro* mineralization defects are present in both male and female *Dock7^em2/em2^* mice. Further studies are necessary to determine whether deletion of *Dock7* exons 3–4 primarily affects late osteoblast differentiation markers, as demonstrated in this study, rather than the early stages of osteoblast differentiation. This work further underscores the critical role of DOCK7 in osteoblast function and mineralization.

In addition to sexually dimorphic changes in cortical bone, total body mass, fat-free mass, aBMD, and aBMC were robustly reduced in *Dock7^em2/em2^* mice, while female mice displayed lower levels of change. Furthermore, *Alpl* expression was only reduced in BMSCs isolated from male *Dock7^em2/em2^ mice*, indicating that deletion of *Dock7* exons 3–4 may present with modified low bone mass phenotypes between male and female mice. While previous studies have reported similarities in the loss of fat-free mass in *Misty* mice, characterization of bone in *Dock7*-mutant mice has primarily been explored in female mice [2, 3]. Our initial observations indicate that BMSCs from male mice express lower levels of DOCK7 protein than those from female mice, suggesting that this differential DOCK7 expression may be one mechanism underlying the observed sex differences in bone. However, we cannot rule out the possibility that the sex differences observed in this study are also linked to the *Dock7* exons 3–4 deletion model itself.

One limitation of this study was that bone microarchitecture was analyzed only at a single time point, 21 weeks of age, where changes in bone microarchitecture between *Dock7^+/+^* and *Dock7^em2/em2^* mice may reflect a combination of bone acquisition and age-related bone loss at the time point analyzed. The current study cannot differentiate whether the observed skeletal phenotype was due to deficits in bone acquisition, enhanced bone loss, or a combination of both. Previous studies suggest that loss of DOCK7 in *Misty* mice primarily affects bone acquisition and possibly early-age-related bone loss, with little impact on long-term age-related bone loss [2, 3]. Therefore, the bone loss in *Dock7^em2/em2^* mice likely reflects a deficiency in bone acquisition. Future experiments should examine different ages to differentiate between these mechanisms.

No studies have indicated that residues from *Dock7* exons 3–4 serve an essential role in protein function. However, the *Dock7^em2/em2^* mouse exhibits a coat color change and low bone mass phenotype similar to the *Misty* model, which shows no detectable protein by Western blot [3]. These results support the hypothesis that *Dock7* exons 3–4 are essential for normal DOCK7 function, making this deletion model a novel tool to study DOCK7 in physiology and cell signaling. Interestingly, the absence of changes in DOCK7 protein levels prior to differentiation suggests the deletion may result in a truncated protein product via an alternative translational start site. While the size of the mutant protein could not be confirmed by Western blot due to the lack of reliable commercial antibodies for mouse DOCK7, SWATH analysis did detect high-confidence peptide fragments in the DHR2 domain. As SWATH failed to detect N-terminal peptides in either genotype, these data do not exclude the presence of a short N-terminal peptide prior to the deletion site.

The presence of these protein products raises the possibility that the *Dock7-em2* allele encodes a nonfunctional, hypomorphic, or dominant-negative protein. A dominant-negative effect is unlikely, however, as *Dock7^+/em2^* mice never exhibit a white belly spot. While we cannot eliminate the possibility that these protein products contribute to the phenotype, as bone was not assessed in heterozygotes, the significant decrease in *Dock7* transcript levels on day 7 of osteogenic differentiation suggests the deletion may instead impair *Dock7* transcript stability during differentiation. Because DOCK7 protein levels were not quantified following osteogenic differentiation, it is also possible that reduced protein levels occur during this stage and contribute to the observed bone phenotype. Further studies are necessary to confirm the functional status of the DOCK7; nonetheless, the phenotypic similarities to *Misty* validate the importance of these exons and indicate the floxed *Dock7-em1* allele will be a valuable tool for future tissue-specific studies.

In summary, the deletion of *Dock7* exons 3–4 replicates key phenotypes observed in *Misty* mice, including trabecular bone deficits and coat color changes. As DOCK7 protein was not detected in *Misty* mice and the low bone mass phenotype was determined to be a result of loss of DOCK7 protein, these data suggest that the *Dock7* exons 3-4 deletion behaves similarly to other Dock7 loss-of-function mutations, supporting its validity as a model for studying the role of DOCK7 in bone physiology. Although the functional status of the truncated DOCK7 protein remains unclear, the phenotypic similarities strongly suggest disrupted normal DOCK7 function. Importantly, the floxed allele design of the *Dock7-em1* model enables future conditional deletion studies to explore cell type-specific roles of DOCK7 in bone development and disease. Further investigation into the functionality of the protein product from the *Dock7-em2* allele will be necessary to understand the molecular mechanisms underlying these phenotypes.

## Acknowledgements

The authors thank Clifford Rosen for providing the *Dock7^em2/em2^* mice to the University of New England. Support from the Center for Cell Signaling and the Center for Pain Research in research development and dissemination of these studies is also gratefully acknowledged. We also thank the University of New England Histology and Imaging Core Facility for technical assistance with von Kossa imaging.

## CRediT

**Talia Lizotte:** Investigation, Data curation, Formal analysis, Writing – original draft, Writing – review & editing.

**Conner Lajoie:** Investigation, Formal analysis, Writing – review & editing.

**Sarah Porter:** Investigation, Data curation, Formal analysis, Writing – original draft, Writing – review & editing.

**Sydney Lang:** Investigation, Writing – review & editing.

**Felix A. Vesper:** Investigation, Writing – review & editing.

**Daniel J. Brooks:** Investigation, Data curation, Formal analysis, Writing – review & editing.

**Carlos Gartner:** Formal analysis, Writing – review & editing.

**Matthew S. Alexander:** Formal analysis, Writing – review & editing.

**Calvin Vary:** Data curation, Methodology, Formal analysis, Writing – original draft, Writing – review & editing.

**Mary L. Bouxsein:** Formal analysis, Writing – review & editing.

**Kathleen A. Becker:** Conceptualization, Investigation, Methodology, Data curation, Funding acquisition, Project administration, Writing – original draft, Writing – review & editing.

## Data availability

Raw data will be deposited in Dryad upon manuscript acceptance. Mass spectrometry (proteomic) data will be deposited into PRoteomics IDEntifications Database, unless otherwise specified.

## Ethics Statement

All animal procedures in this study were conducted in strict accordance with the National Institutes of Health (NIH) *Guide for the Care and Use of Laboratory Animals*. The experimental protocols were reviewed and approved by the Institutional Animal Care and Use Committee (IACUC) at the University of New England (Protocol No. 102219-016, 101922-016) and the MaineHealth Institute for Research (Protocol No. CR1608).

