## Supplemental Figure Legends for "Deletion of *Dock7* Exons 3 and 4 Results in Reduced Trabecular Bone Microarchitecture"

**Supplemental Table 1.** Primers in genotyping and quantitative PCR analysis.

**Supplemental Figure 1. Deletion of *Dock7* exons 3–4 is predicted to generate a premature stop codon in the DOCK7 protein.** (A) Predicted translation of the *Dock7*+ and *Dock7-em2* alleles. Image created with BioRender.com. (B) Examples of genotyping results from $Dock7^{+/+}$, $Dock7^{+/em2}$, and $Dock7^{em2/em2}$ mice.

**Supplemental Figure 2. DOCK7 peptide coverage by mass spectrometry.** DOCK7 peptides were detected by mass spectrometry and used to quantify total DOCK7 levels. High-confidence (green), medium-confidence (yellow), and low-confidence (red) DOCK7 peptides are indicated. The protein sequence corresponding to DOCK7 exons 3–4, the DHR1 domain, and the DHR2 domain is underlined and labeled.

**Supplemental Figure 3. Detection of DOCK7 by mass spectrometry.** (A) Representative DOCK7 peptides. (B) Diagnostic transition intensities for peptide FMDDIAALVSTIAGDVVSR. (C) Ion fragmentation for peptide FMDDIAALVSTIAGDVVSR.

**Supplemental Figure 4. MS/MS collision-induced fragmentation spectra (A) and ion-derived sequence data (B) for indicated diagnostic peptides for BMSCs isolated from $Dock7^{+/+}$ and $Dock7^{em2/em2}$ mice.**

**Supplemental Figure 5. Deletion of *Dock7* exon 3–4 results in no change in *ex vivo* adipose depot weights.** Adipose depots were isolated from both male and female $Dock7^{+/+}$ and $Dock7^{em2/em2}$ mice at 21 weeks of age. The weights of (A) interscapular brown adipose tissue (BAT), (B) inguinal white adipose tissue (WAT), and (C) gonadal WAT were compared by 2-way ANOVA analysis. Points represent each mouse. Statistical significance is indicated: *, $p<0.05$; **, $p<0.01$; ***, $p<0.001$; ****, $p<0.0001$.

**Supplemental Figure 6. Bone density measurements in $Dock7^{em2/em2}$ mice at 20 weeks of age.** Total and femoral areal bone mineral density and content were measured by DXA analysis in male and female

*Dock7*$^{em2/em2}$ mice and compared to *Dock7*$^{+/+}$ control mice. Points represent each mouse. Data was analyzed by 2-way ANOVA and Šídák’s test for post-hoc analysis. Statistical significance is indicated: *, $p<0.05$; **,$p<0.01$; ***, $p<0.001$; ****, $p<0.0001$.
