## Supplemental Table 1 for "Deletion of *Dock7* Exons 3 and 4 Results in Reduced Trabecular Bone Microarchitecture"

| Gene | Application | Forward Primer (5'→3') | Reverse Primer (5'→3') |
|---|---|---|---|
| ***Dock7*+ allele** | **Genotyping** | GCCCAGATCACCCTTGTCTC | TGCTAAGTTTCTAAAAGCTGCCT |
| ***Dock7-em2* allele** | **Genotyping** | GCCCAGATCACCCTTGTCTC | GCAGCCAAGAGGGCAAAAC |
| ***Dock7* exons 11-12** | **qPCR** | GCTTAGAAAGAGACTCCACAGAA | GCCTGCCAACAAGACTAGAAT |
| ***Dock7* exons 34-35** | **qPCR** | GATAAGTCAAGAGCAGAGATAGAAC | TAGCGTGTCTAAGATGATGAGG |
| ***Alpl*** | **qPCR** | GGGACGAATCTCAGGGTACA | AGTAACTGGGGTCTCTCTCTTT |
| ***Bglap*** | **qPCR** | ACCTCACAGATGCCAAGCC | ATCTGGGCTGGGGACTGAG |
| ***Hprt*** | **qPCR** | AAGCCTAAGATGAGCGCAAG | TTACTAGGCAGATGGCCACA |
| ***Runx2*** | **qPCR** | ACCATAACAGTCTTCACAAATCCT | GAGGCGATCAGAGAACAAACTA |
