## Supplementary figures and images for "Deletion of *Dock7* Exons 3 and 4 Results in Reduced Trabecular Bone Microarchitecture"

### Supplemental Figure 1

#### Supplemental Figure 1

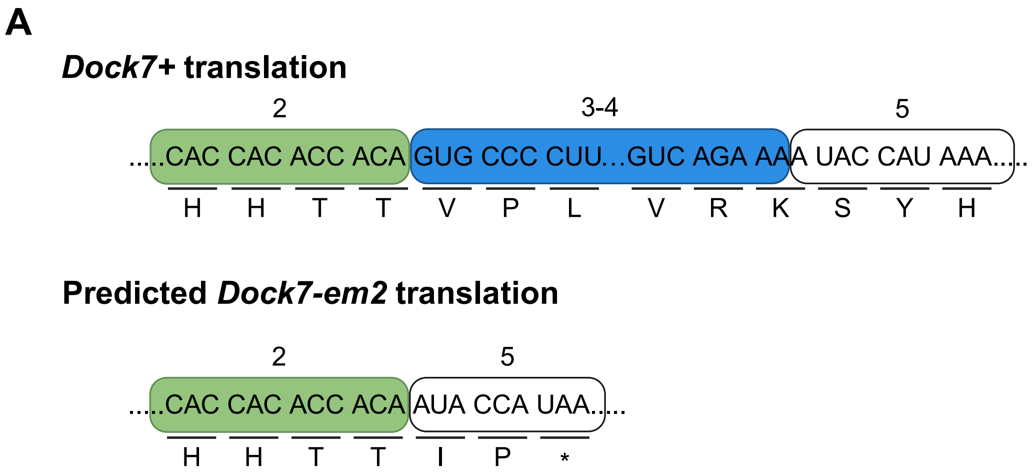


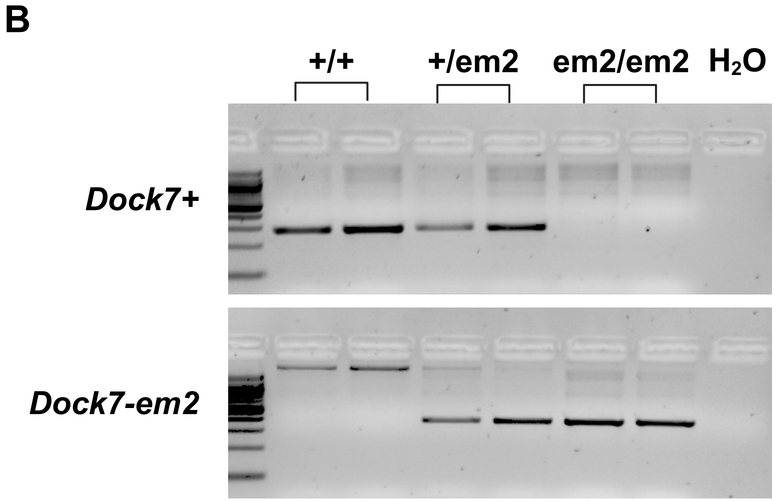

### Supplemental Figure 5

#### Supplemental Figure 5

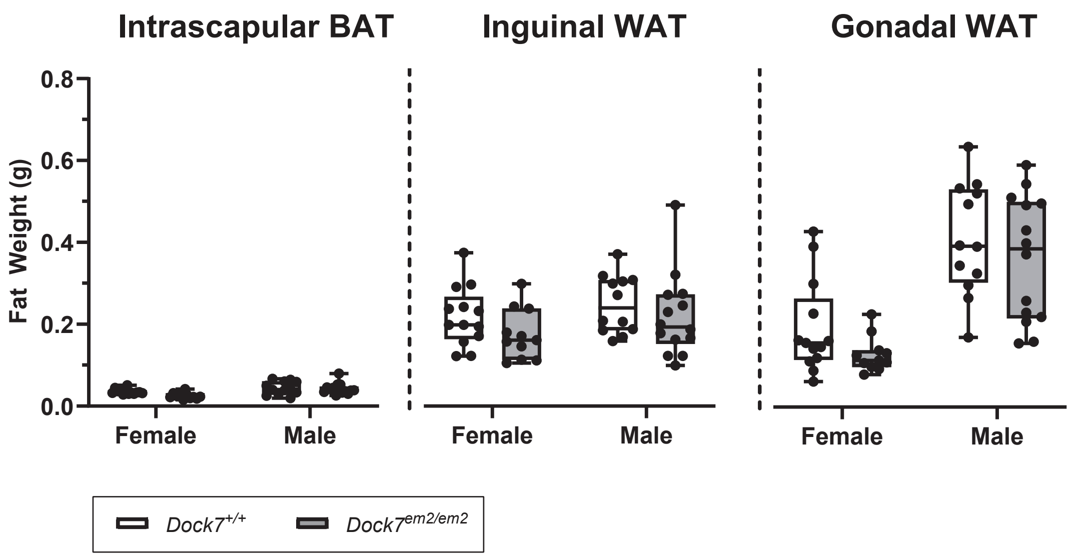

### Supplemental Figure 6

#### Supplemental Figure 6

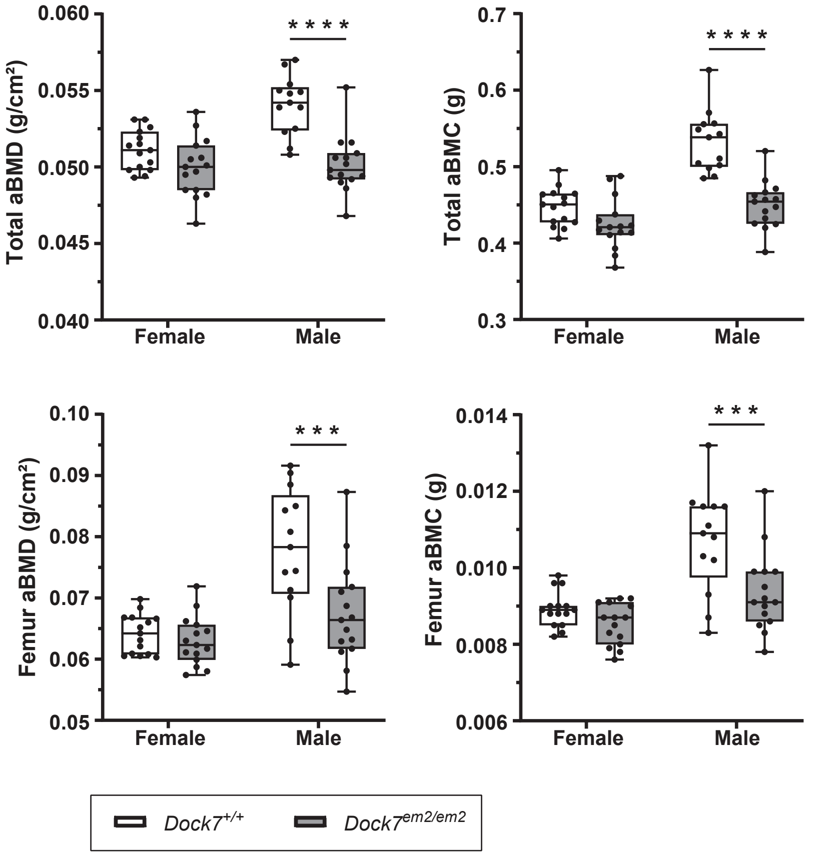
