## Supplemental Figure 2 for "Deletion of *Dock7* Exons 3 and 4 Results in Reduced Trabecular Bone Microarchitecture"

### *Dock7*$^{+/+}$

MAERRAFAQKISRTVAAEVRKQISGQYSGSPQLLKNLNIVGNISHHTTVPLTEAVDPVDLEDYLVTHPLSGDSGPLRDLVEFPPDDIEVVYSPRDCRT
LVSAVPEESEMDPHVRDCIRSYTEDWAVVVRKYHKLGTGFNPNTLDKQKERQKGLPRQVFESDEAPDGSSYQDEQDDLKRRSMSIDDTPRGSW
ACSIFDLKNSLPDALLPNLLDRTPNEEIDHQNDDQRKSNRHKELFALHPSPDEEEPIERLSVPDVPKEHFGQRLLVKCLSLKFEIEIEPIFASLALYDVK
EKKKISENFYFDLNSEQMKGLLRPHVPPAAITTLARSAIFSITYPSQDVFLVIKLEKVLQQGDIGECAEPYMIFKEADATKNKEKLEKLKSQADQFCQR
LGKYRMPFAWTAIHLMNIVSSAGSLERDSTEVEISTGERKGSWSERRNSSLVGRRSLERTTSGDDACNLTSFRPATLTVANFFKQEGDRLSDEDLY
KFLADMRRPSSVLRRLRPITAQLKIDISPAPENPHYCLTPELLQVKLYPDSRVRPTREILEFPARDVYVPNTTYRNLLYIYPQSLNFANRQGSARNITV
KVQFMYGEDPSNAMPVIFGKSSCSEFSKEAYTAVVYHNRSPDFHEEIKVKLPATLTDHHHLLFTFYHVSCQQKQNTPLETPVGYTWIPMLQNGRLK
TGQFCLPVSLEKPPQAYSVLSPEVPLPGMKWVDNHKGVFNVEVVAVSSIHTQDPYLDKFFALVNALDEHMFPVRIGDMRIMENNLESELKSSISAL
NSSQLEPVVRFLHLLLDKLILLVVRPPVIAGQIVNLGQASFEAMASIINRLHKNLEGNHDQHGRNNLLASYIYYVFRLPNTYPNSPSPGPGGLGGSVH
YATMARSAVRPASLNLNRSRSLSNSNPDISGTPTSPDDEVRSIIGSKGLDRSNSWVNTGPKAAPWGSNPSPSAESTQAMDRSCNRMSSHTETSS
FLQTLTGRLPTKKLFHEELALQWVVCSGSVRESALQQAWFFFELMVKSMVHHLYFNDKLDAPRESRFPERFMDDIAALVSTIAGDVVSRFQKDTE
MVERLNTSLAFFLNDLLSVMDRGFVFSLIKSCYKQVSAKLYSLPNPSVLVSLRLDFLRIICSHEHYVTLNLPCSLLTPPASPSPSVSSATSQSSGFSTS
VQDQKIANMFELSLPFRQQHYLAGLVLTELALILDPDAEGLFGLHKKVINMVHNLLSTHDSDPRYSDPQIKARVAMLYLPLIGIIMETVPQLYDFTESH
NQRGRPICIAPDDYDSESGSMISQTVAMAIAGTSVPQLTRPGSFLLTSTSGRQHTTFSAESSRSLLICLLWVLKNADETVLQKWFTDLSVLQLNRLL
DLLYLCVSCFEYKGKKVFERMNSLTFKKSKDMRAKLEEAILGSIGARQEMVRRSRGQLERSPSGSAFGSQENLRWRKDMTHWRQNSEKLDKSRA
EIEHEALIDGNLATEANLIILDTLEIIVQTVSVTESKESILGGVLKVLLQSMACNQSAVYLQHCFATQRALVSKFPELLFEEETEQCADLCLRLLRHCSS
SISTIRSHASASLYLLMRQNFEIGNNFARVKMQVTMSLSSLVGTSQNFNEEFLRRSLKTILTYAEEDLELRETTFPDQVQDLVFNLHMILSDTVKMKE
HQEDPEMLIDLMYRIAKGYQTSPDLRLTWLQNMAGKHSERSNHAEAAQCLVHSAALVAEYLSMLEDRKYLPVGCVTFQNISSNVLEESAVSDDVV
SPDEEGICSGKYFTESGLVGLLEQAAASFSMAGMYEAVNEVYKVLIPIHEANRDAKKLSTIHGKLQEAFSKIVHQDGKRMFGTYFRVGFYGTKFGD
LDEQEFVYKEPAITKLAEISHRLEGFYGERFGEDVLEVIKDSNPVDKCKLDPNKAYIQITYVEPFFDTYEMKDRITYFDKNYNLRRFMYCTPFTLDGR
AHGELHEQFKRKTILTTSHAFPYIKTRVNVTHKEEIILTPIEVAIEDMQKKTQELAFATHQDPADPKMLQMVLQGSVGTTVNQGPLEVAQVFLSEIPGD
PKLFRHHNKLRLCFKDFTKRCEDALRKNKSLIGPDQKEYQRELERNYHRLKEALQPLINRKIPQLYKAVLPVTCHRDSFSRMSLRKMEL

**Exons 3-4**

**DHR-1**

**DHR-2**

### *Dock7*$^{em2/em2}$

MAERRAFAQKISRTVAAEVRKQISGQYSGSPQLLKNLNIVGNISHHTTVPLTEAVDPVDLEDYLVTHPLSGDSGPLRDLVEFPPDDIEVVYSPRDCRT
LVSAVPEESEMDPHVRDCIRSYTEDWAVVVRKYHKLGTGFNPNTLDKQKERQKGLPRQVFESDEAPDGSSYQDEQDDLKRRSMSIDDTPRGSW
ACSIFDLKNSLPDALLPNLLDRTPNEEIDHQNDDQRKSNRHKELFALHPSPDEEEPIERLSVPDVPKEHFGQRLLVKCLSLKFEIEIEPIFASLALYDVK
EKKKISENFYFDLNSEQMKGLLRPHVPPAAITTLARSAIFSITYPSQDVFLVIKLEKVLQQGDIGECAEPYMIFKEADATKNKEKLEKLKSQADQFCQR
LGKYRMPFAWTAIHLMNIVSSAGSLERDSTEVEISTGERKGSWSERRNSSLVGRRSLERTTSGDDACNLTSFRPATLTVANFFKQEGDRLSDEDLY
KFLADMRRPSSVLRRLRPITAQLKIDISPAPENPHYCLTPELLQVKLYPDSRVRPTREILEFPARDVYVPNTTYRNLLYIYPQSLNFANRQGSARNITV
KVQFMYGEDPSNAMPVIFGKSSCSEFSKEAYTAVVYHNRSPDFHEEIKVKLPATLTDHHHLLFTFYHVSCQQKQNTPLETPVGYTWIPMLQNGRLK
TGQFCLPVSLEKPPQAYSVLSPEVPLPGMKWVDNHKGVFNVEVVAVSSIHTQDPYLDKFFALVNALDEHMFPVRIGDMRIMENNLESELKSSISAL
NSSQLEPVVRFLHLLLDKLILLVVRPPVIAGQIVNLGQASFEAMASIINRLHKNLEGNHDQHGRNNLLASYIYYVFRLPNTYPNSPSPGPGGLGGSVH
YATMARSAVRPASLNLNRSRSLSNSNPDISGTPTSPDDEVRSIIGSKGLDRSNSWVNTGPKAAPWGSNPSPSAESTQAMDRSCNRMSSHTETSS
FLQTLTGRLPTKKLFHEELALQWVVCSGSVRESALQQAWFFFELMVKSMVHHLYFNDKLDAPRESRFPERFMDDIAALVSTIAGDVVSRFQKDTE
MVERLNTSLAFFLNDLLSVMDRGFVFSLIKSCYKQVSAKLYSLPNPSVLVSLRLDFLRIICSHEHYVTLNLPCSLLTPPASPSPSVSSATSQSSGFSTS
VQDQKIANMFELSLPFRQQHYLAGLVLTELALILDPDAEGLFGLHKKVINMVHNLLSTHDSDPRYSDPQIKARVAMLYLPLIGIIMETVPQLYDFTESH
NQRGRPICIAPDDYDSESGSMISQTVAMAIAGTSVPQLTRPGSFLLTSTSGRQHTTFSAESSRSLLICLLWVLKNADETVLQKWFTDLSVLQLNRLL
DLLYLCVSCFEYKGKKVFERMNSLTFKKSKDMRAKLEEAILGSIGARQEMVRRSRGQLERSPSGSAFGSQENLRWRKDMTHWRQNSEKLDKSRA
EIEHEALIDGNLATEANLIILDTLEIIVQTVSVTESKESILGGVLKVLLQSMACNQSAVYLQHCFATQRALVSKFPELLFEEETEQCADLCLRLLRHCSS
SISTIRSHASASLYLLMRQNFEIGNNFARVKMQVTMSLSSLVGTSQNFNEEFLRRSLKTILTYAEEDLELRETTFPDQVQDLVFNLHMILSDTVKMKE
HQEDPEMLIDLMYRIAKGYQTSPDLRLTWLQNMAGKHSERSNHAEAAQCLVHSAALVAEYLSMLEDRKYLPVGCVTFQNISSNVLEESAVSDDVV
SPDEEGICSGKYFTESGLVGLLEQAAASFSMAGMYEAVNEVYKVLIPIHEANRDAKKLSTIHGKLQEAFSKIVHQDGKRMFGTYFRVGFYGTKFGD
LDEQEFVYKEPAITKLAEISHRLEGFYGERFGEDVLEVIKDSNPVDKCKLDPNKAYIQITYVEPFFDTYEMKDRITYFDKNYNLRRFMYCTPFTLDGR
AHGELHEQFKRKTILTTSHAFPYIKTRVNVTHKEEIILTPIEVAIEDMQKKTQELAFATHQDPADPKMLQMVLQGSVGTTVNQGPLEVAQVFLSEIPGD
PKLFRHHNKLRLCFKDFTKRCEDALRKNKSLIGPDQKEYQRELERNYHRLKEALQPLINRKIPQLYKAVLPVTCHRDSFSRMSLRKMEL

**Exons 3-4**

**DHR-1**

**DHR-2**
