## Supplemental Figure 3 for "Deletion of *Dock7* Exons 3 and 4 Results in Reduced Trabecular Bone Microarchitecture"

## A

| Peptide Sequence | Charge | Confidence | Intensity | Expected RT | Parent m/z | Observed RT | Score | FDR | % Gaps |
|---|---|---|---|---|---|---|---|---|---|
| IANMFELSLPFR | 2 | 99 | 2098.3 | 63.01 | 719.38 | 62.76 | 5.066 | 0.0 | 19.8 |
| FMDDIAALVSTIAGDVVSR* | 3 | 99 | 6132.82 | 75.35 | 660.68 | 74.79 | 2.58 | 0.0 | 16.7 |
| GSWAC[CAM]SIFDLK | 2 | 99 | 1749.95 | 55.88 | 642.31 | 55.91 | 3.645 | 0.0 | 10.4 |

## B

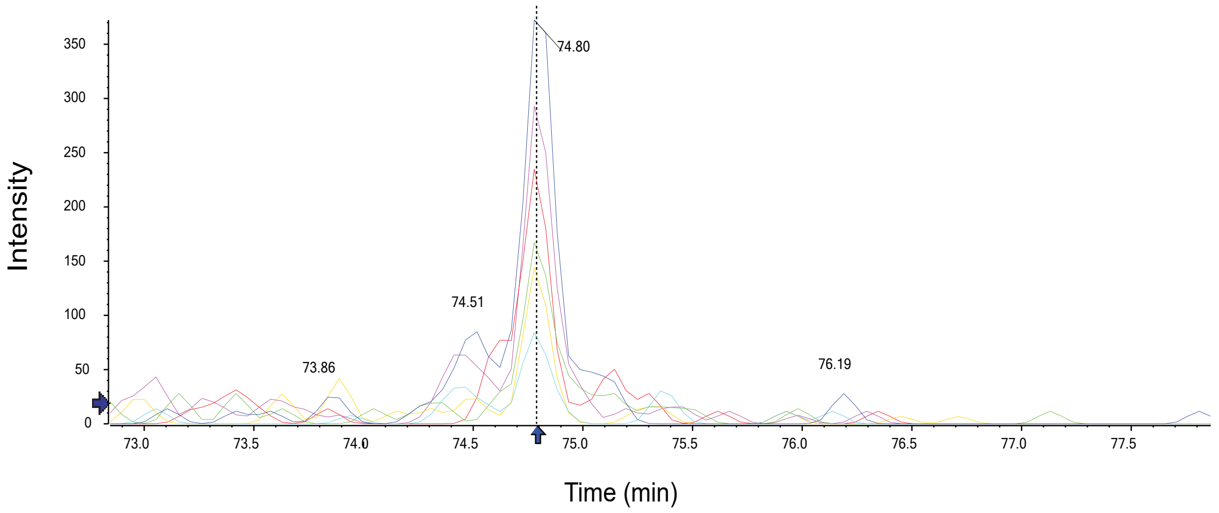


## C

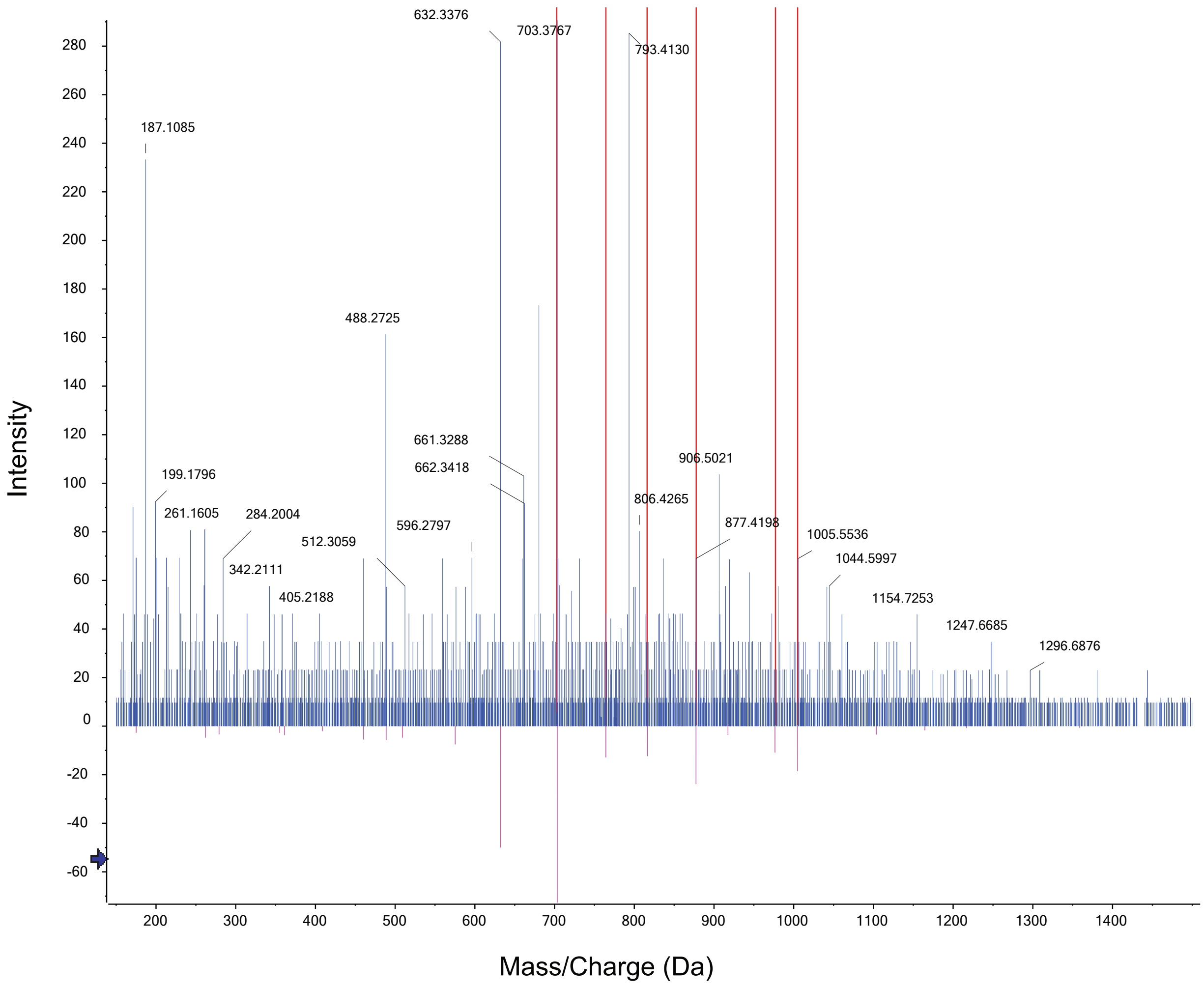
