## Supplemental Figure 4 for "Deletion of *Dock7* Exons 3 and 4 Results in Reduced Trabecular Bone Microarchitecture"

**A**

***Dock7$^{+/+}$***

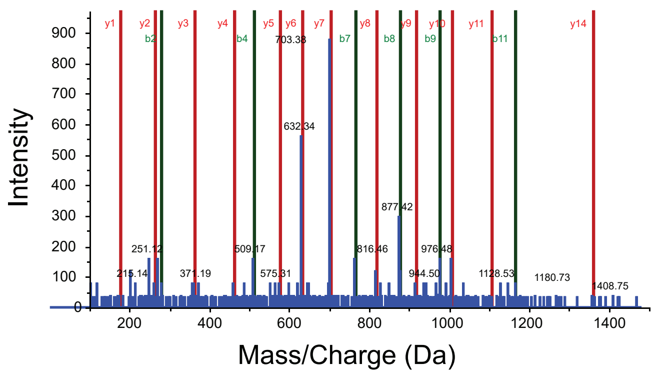


***Dock7$^{em2/em2}$***

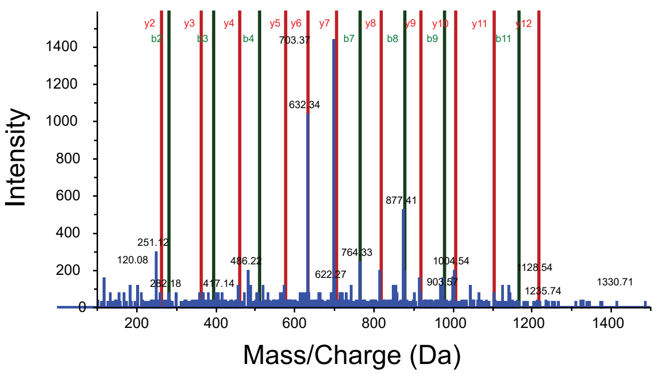


**B**

***Dock7$^{+/+}$***

| Residue | b | y |
|---|---|---|
| F | 148.0757 | 1980.0107 |
| M | 279.1162 | 1832.9422 |
| D | 394.1431 | 1701.9018 |
| D | 509.1701 | 1586.8748 |
| I | 622.2541 | 1471.8479 |
| A | 693.2912 | 1358.7638 |
| A | 764.3284 | 1287.7267 |
| L | 877.4124 | 1216.6896 |
| V | 976.4808 | 1103.6055 |
| S | 1063.5129 | 1004.5371 |
| T | 1164.5605 | 917.5051 |
| I | 1277.6446 | 816.4574 |
| A | 1348.6817 | 703.3733 |
| G | 1405.7032 | 632.3362 |
| D | 1520.7301 | 575.3148 |

***Dock7$^{em2/em2}$***

| Residue | b | y |
|---|---|---|
| F | 148.0757 | 1980.0107 |
| M | 279.1162 | 1832.9422 |
| D | 394.1431 | 1701.9018 |
| D | 509.1701 | 1586.8748 |
| I | 622.2541 | 1471.8479 |
| A | 693.2912 | 1358.7638 |
| A | 764.3284 | 1287.7267 |
| L | 877.4124 | 1216.6896 |
| V | 976.4808 | 1103.6055 |
| S | 1063.5129 | 1004.5371 |
| T | 1164.5605 | 917.5051 |
| I | 1277.6446 | 816.4574 |
| A | 1348.6817 | 703.3733 |
| G | 1405.7032 | 632.3362 |
| D | 1520.7301 | 575.3148 |
